# The cochlear hook region detects harmonics beyond the canonical hearing range

**DOI:** 10.1101/2024.04.14.589450

**Authors:** Kazuhiro Horii, Bakushi Ogawa, Noriko Nagase, Iori Morimoto, Chikara Abe, Takenori Ogawa, Samuel Choi, Fumiaki Nin

## Abstract

Ultrasound, or sound at frequencies exceeding the conventional range of human hearing, is not only audible to mice, microbats, and dolphins, but also creates an auditory sensation when delivered through bone conduction in humans. Although ultrasound is utilized for brain activation and in hearing aids, the physiological mechanism of ultrasonic hearing remains unknown. In guinea pigs, we found that ultrasound above the hearing range delivered through ossicles of the middle ear evokes an auditory brainstem response and a mechano-electrical transduction current through hair cells, as shown by the local field potential called the cochlear microphonic potential (CM). The CM synchronizes with ultrasound, and like the response to audible sounds is actively and nonlinearly amplified. In vivo optical nano-vibration analysis revealed that the sensory epithelium in the hook region, the basal extreme of the cochlear turns, resonates in response both to ultrasound within the hearing range and to harmonics beyond the hearing range. The results indicate that hair cells can respond to stimulation at the optimal frequency and its harmonics, and the hook region detects ultrasound stimuli with frequencies more than two octaves higher than the upper limit of the ordinary hearing range.

## Introduction

Ultrasound refers to those frequencies that exceed the upper limit of audibility in humans, which lies at approximately 20 kHz. In other species, such as in mice and bats, the cochlea has instead evolved to make use of much higher frequencies for communication. Moreover, the capacity of ultrasound to activate brain circuits noninvasively has positioned it as a promising candidate for neuromodulation techniques (Fisher & Gumenchuk, 2018; Nitsche et al., 2008). Audible sound elicits a wave that traverses the sensory epithelium from the base of the cochlea towards the apex and whose amplitude peaks at a frequency-dependent location. High frequencies evoke a response at the cochlear basal end, whereas lower frequencies stimulate the most apical edge (Bekesy, 1960; Gelfand, 2010; Robles & Ruggero, 2001; Ulfendahl, 1997). This spatial arrangement of best frequencies, called tonotopy, formally demarcates the hearing range in various animal species (Liberman, 1982; Manley, 2017; Ruggero & Temchin, 2002). In humans, ultrasonic frequencies by definition fall outside the range of tonotopically arranged frequencies, yet they can still be perceived when presented through bone conduction (Corso, 1963; Gavreau, 1948). This extraordinary feature, known as ultrasonic hearing, is utilized for hearing aids and tinnitus sound therapy (Koizumi et al., 2014; Nishimura et al., 2011).

Although the neural circuit including auditory cortex can be activated by ultrasonic stimuli (Fisher & Gumenchuk, 2018), (Guo et al., 2018; Hosoi, Imaizumi, Sakaguchi, Tonoike, & Murata, 1998; Sato, Shapiro, & Tsao, 2018), the physiological basis of both ultrasonic hearing and ultrasound-induced brain activation remains unclear. The fact that neurons and ion channels can be stimulated by ultrasound has led some to propose its potential for directly activating the auditory cortex (Hoffman et al., 2022; Kubanek et al., 2016; Tyler et al., 2008). At the same time, transection of the auditory nerves or removal of cochlear fluids eliminates the ultrasound-induced neural activities (Guo et al., 2018), and ultrasonic-induced sensations can be masked by audible sounds (Nishimura, Nakagawa, Sakaguchi, & Hosoi, 2003; Nishimura et al., 2011). Collectively, these studies imply that the cochlea plays a significant role in ultrasonic hearing.

During air conduction, sound travels to the cochlea by way of the tympanic membrane and middle ear ossicles. In bone conduction that is increasingly employed in headphones that bypass the middle ear, sounds instead engage the cochlea directly through the temporal bone (Bekesy, 1960; Gelfand, 2010). Bone conduction is thought to be indispensable in ultrasonic hearing. Although theoretical studies have demonstrated that both air- and bone-conducted sounds evoke similar nanoscale vibrations in the cochlear sensory epithelium (Reichenbach & Hudspeth, 2010; Tchumatchenko & Reichenbach, 2014), it remains unclear why only bone-conducted ultrasound can be audible. From the mechanical perspective, there are two possibilities: First, the bone may mechanically convert ultrasound into audible sounds (Dieroff & Ertel, 1975; Haeff & Knox, 1963). Second, bone-conducted ultrasound might bypass the tympanic membrane’s strong low-pass filtering effect (Cooper & Rhode, 1992) and directly reach the cochlea. However, no detailed studies have been performed *in vivo* to test these assumptions.

In this study, we measured auditory brainstem response (ABR) and cochlear microphonic potentials (CM)—a proxy of the mechanoelectrical transduction (MET) current through stimulated hair cells—in guinea pigs in order to establish the range of detectable frequencies in different types of sound conduction. Ultrasound above the animal’s hearing range elicited ABR and CM not only by bone conduction but also by stimulation through the malleus-incus complex with a tapered stainless rod. Based on the tonotopic arrangement of the sensory epithelia, we focused on the cochlear hook region—the extreme base of the cochlea—as the probable location of ultrasound detection. We optically confirmed that the hook region receives ultrasound stimuli both at the optimal frequency and at its harmonics, exceeding the upper limit of the conventional hearing range.

## Results

### Auditory brainstem response under ultrasonic stimulation through middle-ear ossicles

The hearing range is conventionally defined as the band of frequencies over which an animal responds to air-conducted sounds (**Figure 1A**). We first performed control experiments, meant to confirm the hearing range of the guinea pigs. We measured the ABR in vivo by exposing the cochlea to air-conducted sound covering frequencies ranging from 10 kHz to 80 kHz. The minimum threshold was 30 dB at a frequency of 16 kHz. As the stimulus frequency increased to 40 kHz, which is near the upper limit of the hearing range, the thresholds increased to 65 dB. During 80 kHz stimulation at 80 dB, ABR was not observed (**Figure 1B**). This result, which is consistent with that of a previous study (Abaamrane, Raffin, Gal, Avan, & Sendowski, 2009), indicates that the conventional hearing range of the guinea pig extends slightly above 40 kHz, and that stimuli of substantially greater frequencies constitute ultrasound for that species.

**Figure 1.**
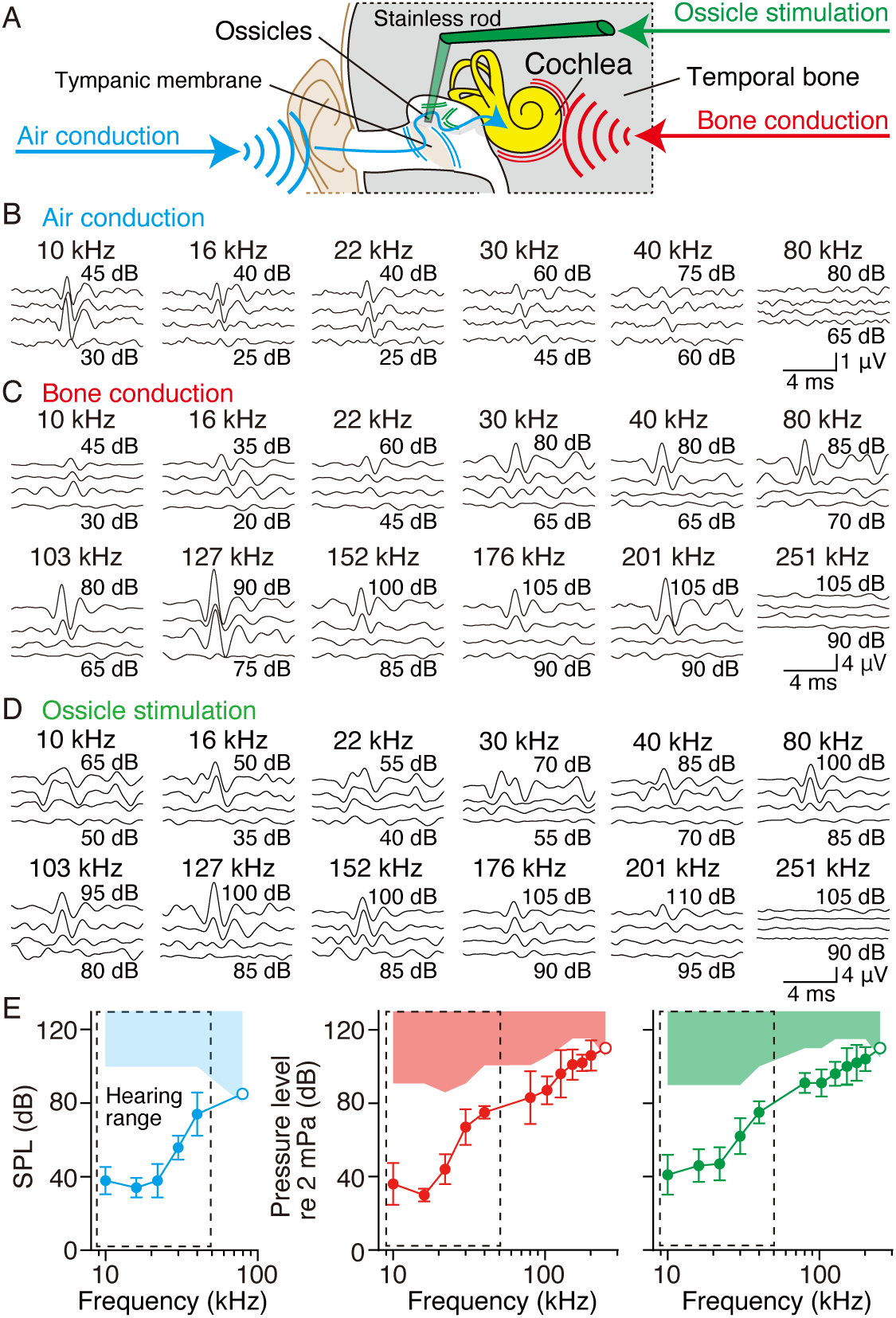
ABR signals and thresholds in air and bone conduction. (A) Schematic diagrams of air and bone conduction and ossicle stimulation. Sound vibrations pass from the external ear canal to the cochlea by the tympanic membrane in air conduction (blue arrow), whereas bone-conducted vibration directly reaches the cochlea through the temporal bone (red arrow). In ossicle stimulation (green arrow), vibration is applied through the malleus-incus complex in the air-conduction pathway. (B-D) Representative examples of ABR signals under air- or bone-conducted stimulation and ossicle stimulation in guinea pigs. The results are plotted in 5 dB decrements. (E) Grouped ABR thresholds for various types of stimulation in guinea pigs. The left, middle, and right panels show the air conduction (blue, n = 5), bone conduction (red, n = 5), and ossicle stimulation (green, n = 5) data, respectively. The black dashed rectangles show the hearing range; the blue, red, and green shaded areas indicate inapplicable pressure levels for each stimulation. ABR thresholds under air and bone-conducted stimulations showed almost the same thresholds at audible frequencies, but ABR thresholds were observed at ultrasound frequencies in bone conduction and ossicle stimulation. For each stimulation, data are shown as Mean ± SD. The open circles indicate that the ABR thresholds are out of range.

We next measured the ABR during bone-conducted sound stimulation through the temporal bone (**Figure 1A**). Within the conventional hearing range, ABR thresholds were similar to those observed in air conduction (**Figure 1C**). For frequencies exceeding the conventional hearing range, the ABR was not detected at intensities less than 70 dB, but became prominent at intensities exceeding 70 dB (**Figure 1C**). Thresholds increased monotonically within frequencies from 80 kHz to 201 kHz; no ABR was recorded at frequencies above 251 kHz with 105 dB stimulation.

During air conduction, a pressure wave is transmitted from the tympanic membrane to the cochlea through the middle-ear ossicles. To avoid the low-pass filtering effect of the tympanic membrane (Cooper & Rhode, 1992), we directly applied stimulation to the malleus-incus complex (**Figure 1A**). Although the stimulation was applied along the air conduction pathway, ABR thresholds were similar to those in bone conduction (**Figure 1D** and **E**), suggesting that otherwise inaudible ultrasounds can also elicit ultrasonic hearing when delivered through direct stimulation of the ossicles. As a result, the detectable frequency ranges in both ossicle stimulation and bone conduction are broader than the ordinary hearing range.

The reference pressures in bone conduction and ossicle stimulation differed from those of air conduction. The pressure was calibrated with a microphone for air conduction and by means of a hydrophone in the cases of bone conduction and ossicle stimulation. To roughly compensate for the threshold difference between these modes of conduction within the hearing range, we defined the standard pressure in bone conduction and ossicle stimulation as 2 mPa and show pressure levels in decibels throughout this study. In addition, although audible sounds greater than 105 dB generally induce acoustic trauma when delivered through air conduction, they did not elevate the threshold when delivered through bone conduction (**Figure S1A-C**).

### Cochlear microphonic potential during ultrasonic stimulation

Because inaudible ultrasonic stimuli evoke ABR responses when delivered through both bone conduction and direct ossicle stimulation, we hypothesized that the hair cells of the cochlea must be capable of transducing sound stimuli at much higher frequencies that previously thought impossible. A standard criterion for evaluating the proper functioning of hair cells is their ability to generate MET currents. To assess the integrity of this mechanism, we recorded the CM using a silver electrode placed within the middle ear cavity (**Figure 2A**). The CM is the alternating-current component of the local field potential (LFP) generated by excitable cells in the cochlea (Gelfand, 2010; Tasaki, 1952; Wever & Bray, 1930). Although CM responses to bone-conducted ultrasonic stimulation in guinea pigs have been documented for stimuli at 98.8 kHz, its occurrences at other frequencies remains unexplored (Ohyama, Kusakari, & Kawamoto, 1985).

**Figure 2.**
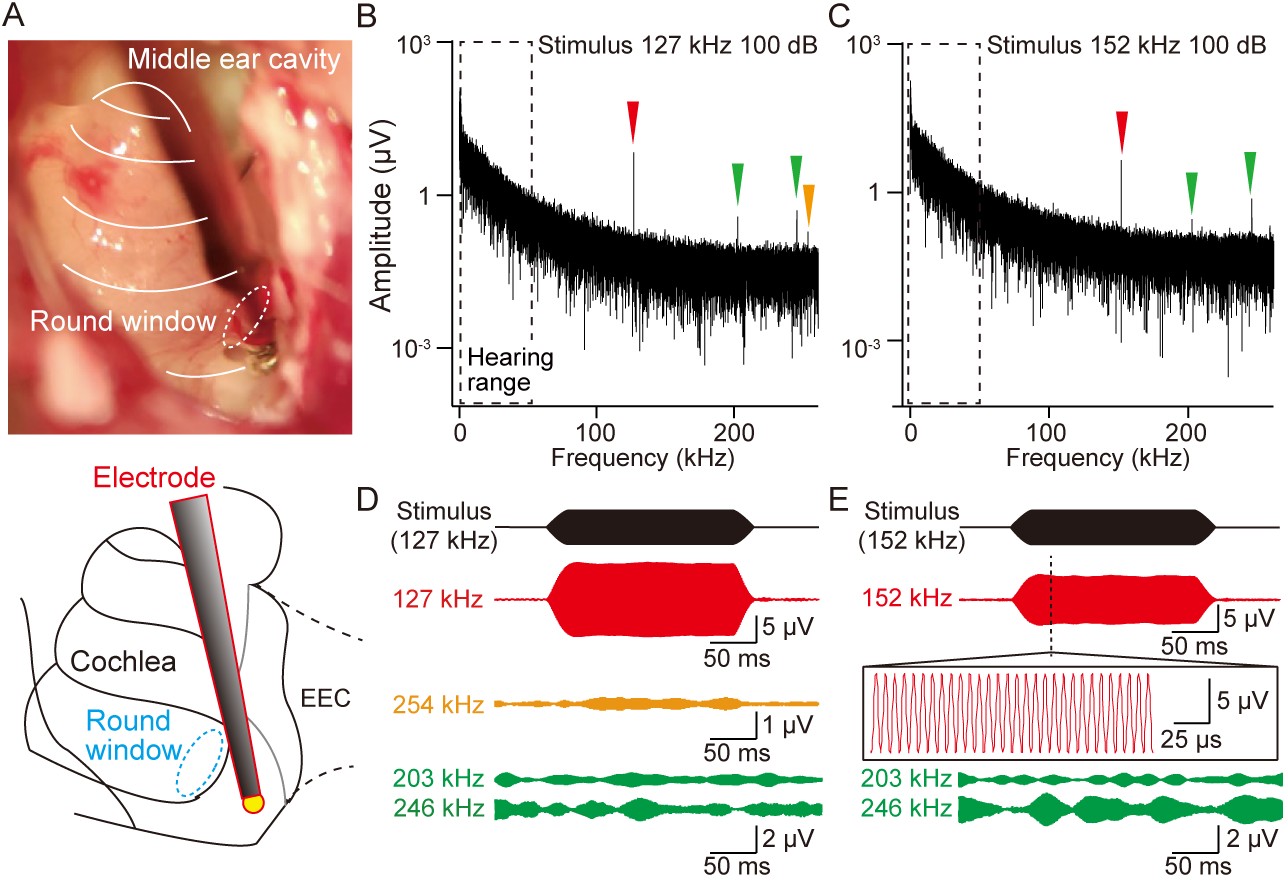
Recording of the local field potentials (LFP) in a guinea pig. (A) A view of the middle ear cavity with the recording electrode in place. The upper panel shows a photo of the middle ear cavity, and the lower panel shows its schematic representation. The tip of the recording electrode was placed on the middle ear mucosa near the round window. EEC, external ear canal. (B, C) Fourier amplitude spectrum of the LFP during stimulation at127 kHz and 152 kHz at 100 dB. The arrowheads demark the characteristic frequency peaks of the persistent electrical signals in the experiments. The red and orange filled arrowheads indicate stimulus frequencies and harmonics, respectively. The black dashed rectangles delimit the hearing range of the animal. (D) LFP waveforms at 127 kHz (red), 254 kHz (orange), 203 kHz, and 246 kHz (green) with a stimulus waveform (black) in (B). (E) LFP waveforms at 152 kHz (red), 203 kHz, and 246 kHz (green) in C. A zoomed view of the LFP at 152 kHz under steady-state stimulation is shown in the insert. Signals at 127 kHz, 254 kHz, and 152 kHz are synchronized with the stimulus, whereas those at 203 kHz and 246 kHz are asynchronous.

Prior to analyzing the CM responses to various stimuli, we evaluated the frequency distribution of the LFP during stimulation through the temporal bone. **Figures 2B** and **C** display the fast Fourier transform (FFT) of the voltage amplitude measured from the round window membrane. First, we delivered a 127 kHz stimulus of 100 dB to the temporal bone of the animal. Significant peaks were visible at 127 kHz, 254 kHz, 203 kHz and 246 kHz (**Figure 2B**). **Figures 2D** and **E** exhibit the LFP waveforms with a stimulus waveform at the peaks. Waveforms at 127 kHz and 254 kHz, a harmonic of 127 kHz, were synchronized with a stimulus waveform. When the applied frequency was changed from 127 kHz to 152 kHz, the peak was shifted to 152 kHz, but the waveform remained synchronous (**Figure 2E**). In contrast, the signals at 203 kHz and 246 kHz under stimulations both of 127 kHz and of 152 kHz were asynchronous (**Figure 2D** and Based on the frequency dependence (**Figure 2B** and **C**), we defined the waveforms at peaks of 127 kHz and 152 kHz as ultrasonic CMs. Conversely, the asynchrony to the stimulus suggests that the waveforms at 203 kHz and 246 kHz represented physical or electrical resonances in the recording. Notably, LFP signals within the hearing range were not detected (**Figure 2B** and **C**). In additional, harmonics of the ultrasonic CM were occasionally observed during the intense stimulation through bone conduction (**Figure S2A**), whereas no harmonics were detected during ossicle stimulation (**Figure S2B**). These results indicate that objects in the bone conduction pathway do not mechanically convert ultrasonic vibration into audible sounds.

In normal hearing, a healthy cochlea nonlinearly receives weak inputs while progressively reducing the enhancement of stronger stimuli. This level-dependent reception is known as compressive nonlinearity based on an active process of the cochlea (Hudspeth, 2008), and the sound-evoked CM exhibits its signature (Honrubia & Ward, 1968). To determine whether a similar pattern occurs in the ultrasonic CM, we analyzed the level function of the amplitude. The root-mean-square voltage was acquired while stimulating with a sinusoidal stimulus; the frequency and pressure level ranged from 80 kHz to 251 kHz and from 70 dB to 100 dB, respectively (**Figure 3A, S3A and B**). Under control conditions, the amplitude displayed clear compressive nonlinearity at frequencies of 80, 103, and 127 kHz. In contrast, the amplitude showed slight nonlinearity at the frequencies of 152, 176, and 201 kHz (**Figure 3B** and **C**). These values were significantly higher than those obtained under postmortem conditions, as shown by the limit of detection (LOD). These results indicate that the cochlea houses ultrasound-sensitive hair cells whose response is actively and nonlinearly amplified. In addition, the amplitudes under normal stimulation and LOD were almost the same under a high-frequency stimulus of 251 kHz (**Figure S3A and B**). The upper frequency limit of the CM response was almost identical to that of the bone-conducted hearing range determined by ABR (**Figure S1E**). These tendencies were also observed in recordings during ultrasonic stimulation by means of the ossicles (**Figure S3C and D**).

**Figure 3.**
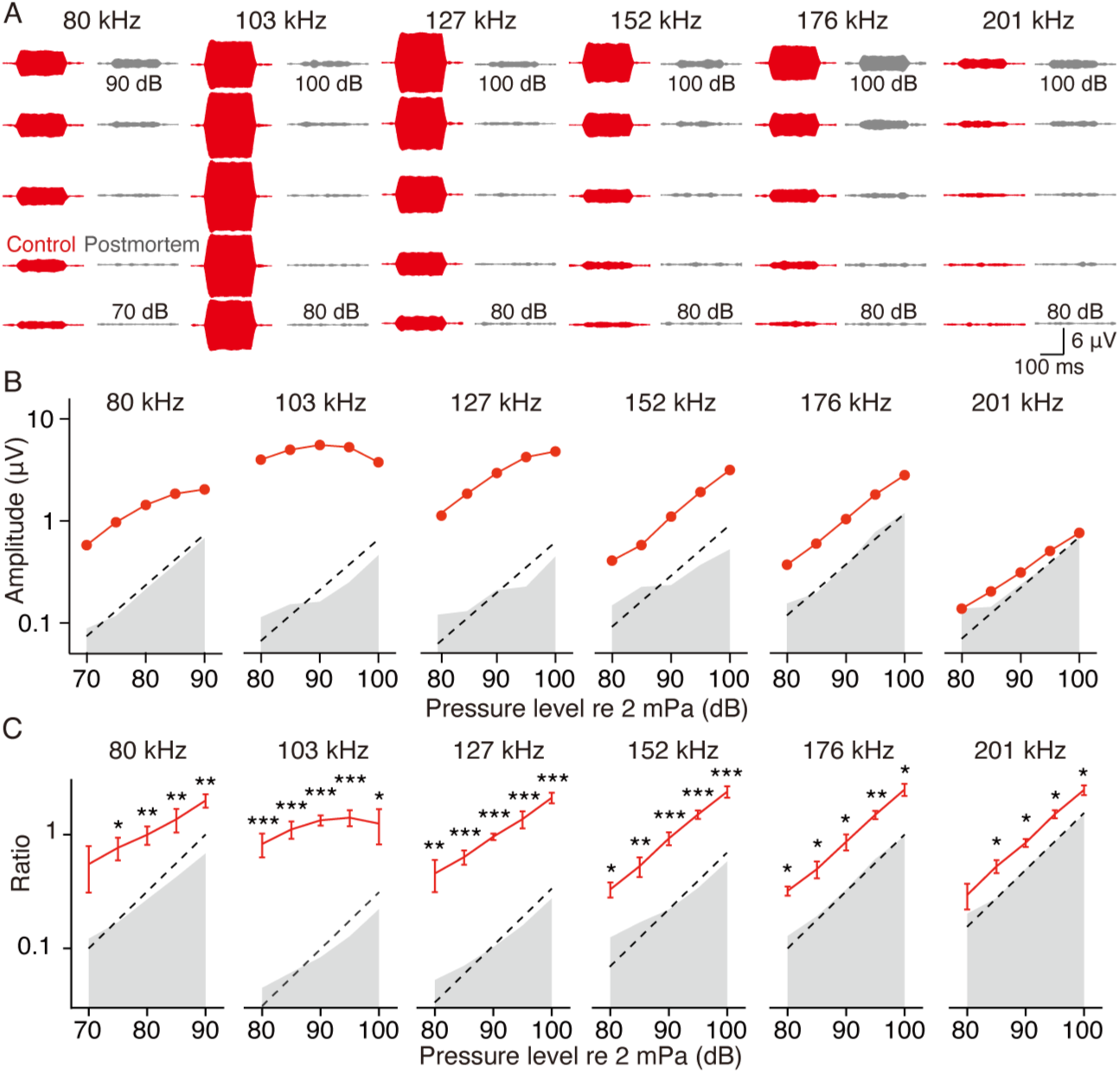
Recording of the cochlear microphonic potentials (CM) in a guinea pig. (A) Representative trace of stimulation-evoked CM waveforms in a guinea pig. The results are plotted in 5 dB decrements. The left and right panels for each stimulus frequency show the control and postmortem conditions, respectively. (B) Summary profile of CM amplitude and its nonlinear response in the guinea pig. The level function relates to the amplitude of CM responses in the cochlea. Control data (red) during 80, 103, and 127 kHz stimulation demonstrate strong compressive nonlinearity at higher pressure levels, whereas those during 152, 176, and 201 kHz exhibit slightly nonlinear behavior. The thin dashed lines mark a linear relationship between the pressure level and CM amplitude. Because an actuator elicits electrical artifacts synchronized to the applied voltage frequency during stimulation, we defined the average amplitude of the recorded artifacts under postmortem as the limit of detection (gray shaded area). The high-resolution ultrasonic waveforms with a sampling frequency of 2.4 MHz are downsampled to 24 kHz throughout this study. (C) Grouped CM amplitude in five guinea pigs (n = 5). The solid line and error bars indicate the averages and standard deviations, respectively, for the control conditions, whereas gray shadings indicate the averages for postmortem. The responses were normalized to the average values of the recorded voltages under control conditions and are shown as ratios. *, ** and *** : significant differences (*p* < 0.32, *p* < 0.05, and *p* < 0.01)

### Location of ultrasound detection in the cochlea

The hook region, which curves in the opposite direction to that of the cochlear spiral, represents the base of the cochlea. In the hook region of the cochlea, high-frequency sounds near the upper limit of the hearing range are thought to elicit a peak wave of epithelial motion (Cooper & Rhode, 1992; Gelfand, 2010; Liberman, 1982; Manley, 2017; Robles & Ruggero, 2001; Ruggero & Temchin, 2002; Ulfendahl, 1997). This frequency-dependent spatial arrangement underlies cochlear frequency tuning. We hypothesized that the hook region can transduce ultrasound at frequencies exceeding 100 kHz into nanoscale vibrations in the sensory epithelium.

As the initial step in each experiment, we focused the OCT beam onto the epithelium through the round window, and positioned the focus of the objective lens at the depth of the reticular lamina (RL) (**Figure 4A**). The depth of focus was 1.7 μm. The epithelium extends from the RL, which includes the apical surfaces of outer hair cells (OHCs), to the basilar membrane (**Figure 4B**). On a two-dimensional cross-sectional image, the RL was visualized as the apical edge of the organ of Corti (**Figure 4C**). A representative signal on the pathway of the beam passing through OHCs is shown in **Figure 4B** and **C**. The intensities of RL exceeded the background signal in the endolymph by more than 3 dB (**Figure 4D**). To measure the vibration profiles of the epithelium by OCT vibrometry, we subsequently administered an ultrasonic stimulus between 35 kHz and 130 kHz at pressure intensity levels of 35 dB, 45 dB, and 55 dB. **Figure 4E** shows the amplitude of vibrations at the RL at the stimulus frequency (f_1_), whereas **Figure 4F** shows those at the 2nd and 3rd harmonics of f_1_ at 55 dB. **Figure 4G** exhibits the frequency distribution of the amplitudes. The amplitudes were recorded as the maximal values around the peak signal intensities. In general, the stimulus frequency that maximally vibrates the epithelium with a low pressure at the recording point represents the best frequency (BF) (Ota et al., 2020; Strimbu & Olson, 2022). During moderate stimulation at 55 dB, the vibration amplitudes were highest at approximately 45 kHz, whereas no significant amplitudes were observed at 35 kHz or at frequencies exceeding 60 kHz. Based on the frequency response to ultrasound, the epithelium resonated at frequencies of approximately 45 kHz within the hearing range. Therefore, we categorized the 45 kHz stimuli as near-BF, those at 35 kHz as sub-BF, and those over 60 kHz as supra-BF in this trial. In near-BF stimulations, the amplitude at 55 dB was less than 3.1 times that at 45 dB, and harmonics were significant (**Figure 4G**). These phenomena were similar to those at the apical and basal turns in previous studies (Dewey, Altoe, Shera, Applegate, & Oghalai, 2021) (Cooper, 1998). In contrast, no harmonic vibration was observed for sub- and supra-BF stimulation (**Figure 4G**). The vibrational phases in the epithelium displayed little level dependence (**Figure 4H),** whereas the phases near the RL led those in the OHC body in the near-BF (**Figure S4**).

**Figure 4.**
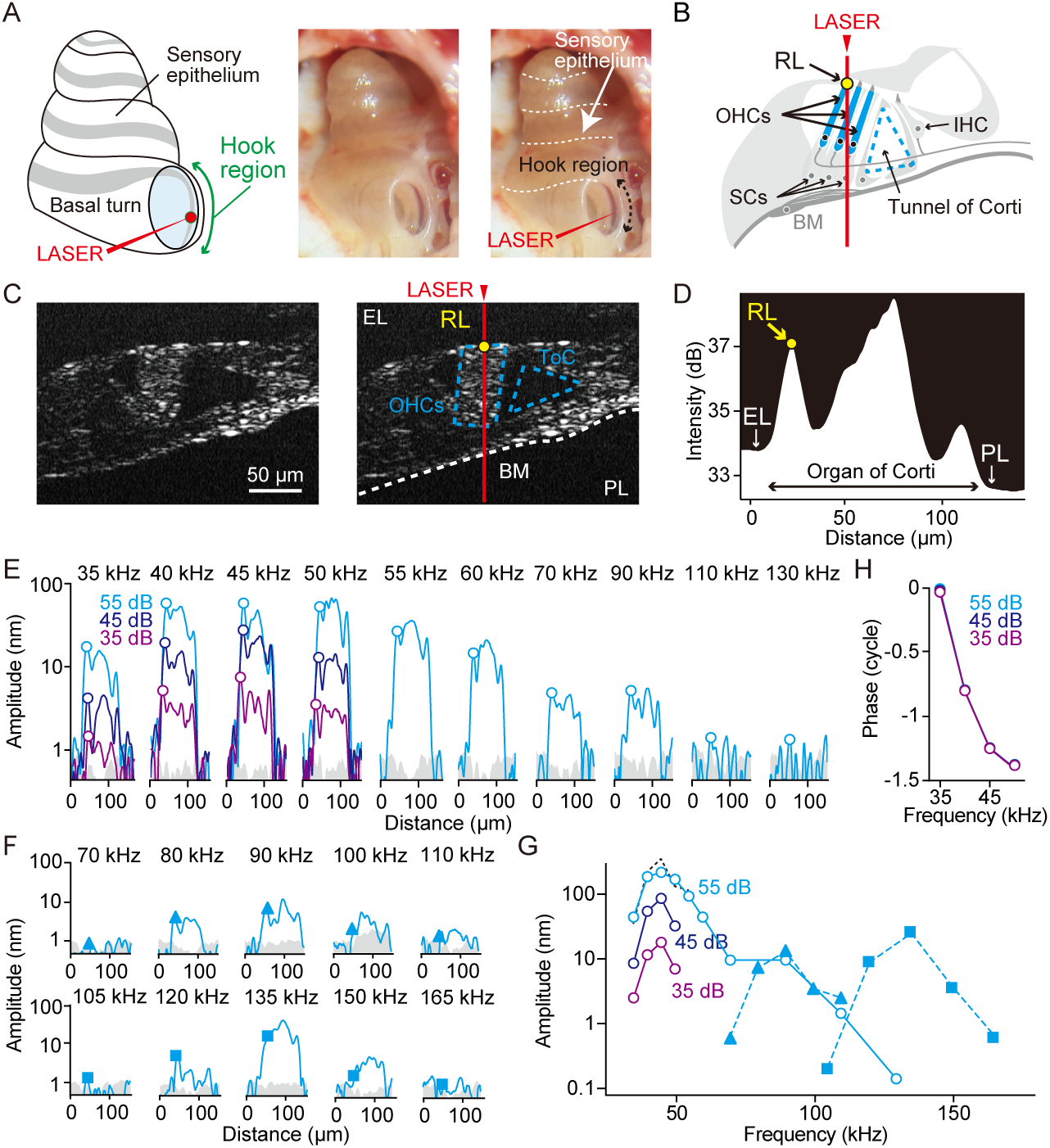
Optical coherence tomography (OCT) vibrometry analyses of the organ of Corti in the cochlear hook region during ultrasound stimulation within the hearing rage. (A) A schematic drawing of the cochlea and images of the sensory epithelium through the round window during a recording with and without landmarks. (B) Schematics of a cochlear partition. RL, reticular lamina, IHC, inner hair cell, OHCs, outer hair cells, SCs, supporting cells, BM, basilar membrane. (C) OCT image of the epithelium in the hook region with and without indicative landmarks. A cluster of OHCs and the tunnel of Corti (ToC) are highlighted with dashed blue lines. A yellow filled circle indicates the representative points on the RL. The red arrowhead indicates the pathway of the OCT beam and the selected line shown in (D). (D) Reflection intensity along the selected line plotted on a logarithmic scale. The top of the cochlear partition lies to the right and the bidirectional arrow represents the cochlear partition. The reflection intensities of the RL are shown as a yellow circle. PL, perilymph. EL, endolymph. (E) Vibration amplitudes in response to stimulation from 35 kHz to 130 kHz at 35, 45 and 55 dB under physiological conditions. The open circles indicate the amplitudes at the RL. In this and subsequent illustrations, the results of different stimulations are plotted in a chromatic sequence. The gray shaded area shown as the limit of detection corresponds to the amplitude measured during 55 dB stimulation. (F) Vibration amplitudes of the 2nd and 3rd harmonics in response to stimulation from 35 kHz to 55 kHz with an intensity of 55 dB. (G) Frequency distribution of the vibration amplitudes for the RL. The results of stimulations at the fundamental frequency and harmonics are plotted as open circles, filled triangles, and filled squares, respectively. The dashed black line shows the calculated linear increment in the amplitude for the 45 dB stimulus. (H) Frequency distribution of the phase for the RL in reference to the phase of the stimulation.

Because harmonic vibrations were detected during near-BF stimulation, the hook region resonates physically with harmonics as well as the fundamental frequency. We therefore defined the double and triple BF as the 2nd and 3rd harmonic BF (2hBF, 3hBF), respectively. The maximum amplitude of the 3rd harmonic was greater than that of the 2nd harmonic (**Figure 4G**); thus we further delivered an ultrasonic stimulus at frequencies greater than 116 kHz and pressure intensities of 70 dB, 75 dB, 80 dB, and 85 dB. **Figure 5A** shows the frequency distributions of the amplitudes. During the stimulation, the vibration amplitudes were largest near 122 kHz, whereas the amplitudes were not significant at 116 kHz. Therefore, we noted the 122 kHz stimuli as near-3hBF, and those below 116 kHz as sub-3hBF. In near-3hBF stimulations, the amplitude at 85 dB was smaller than 3.1 times that at 75 dB for RL (**Figure 5B**). This indicates that the amplitudes exhibit compressive nonlinearity. In contrast, the amplitudes increased more linearly for sub-3hBF stimulation. Although guinea pigs do not seem to hear these ultrasounds under ordinary conditions, the result was comparable to the vibration response in the epithelium of other cochlear turns during stimulation with audible sounds (F. Nin, Choi, Ota, Qi, & Hibino, 2021; Strimbu & Olson, 2022). The vibrational phases for RL in reference to the stimulus signal and the relative phase distribution inside the OHCs are shown in **Figure 5C** and **D**, respectively. There was little variation in phase for RL across stimulus levels (**Figure 5C**). However, phases near the RL led those at OHC’s body (**Figure 5D**), and the rates of the phase change in near-3hBF stimulation were significantly higher than those in sub-3hBF stimulation at lower stimulus levels (**Figure 5E**). These trends in amplitudes and phases were confirmed in five experiments (**Figure 5F** and **G**).

**Figure 5.**
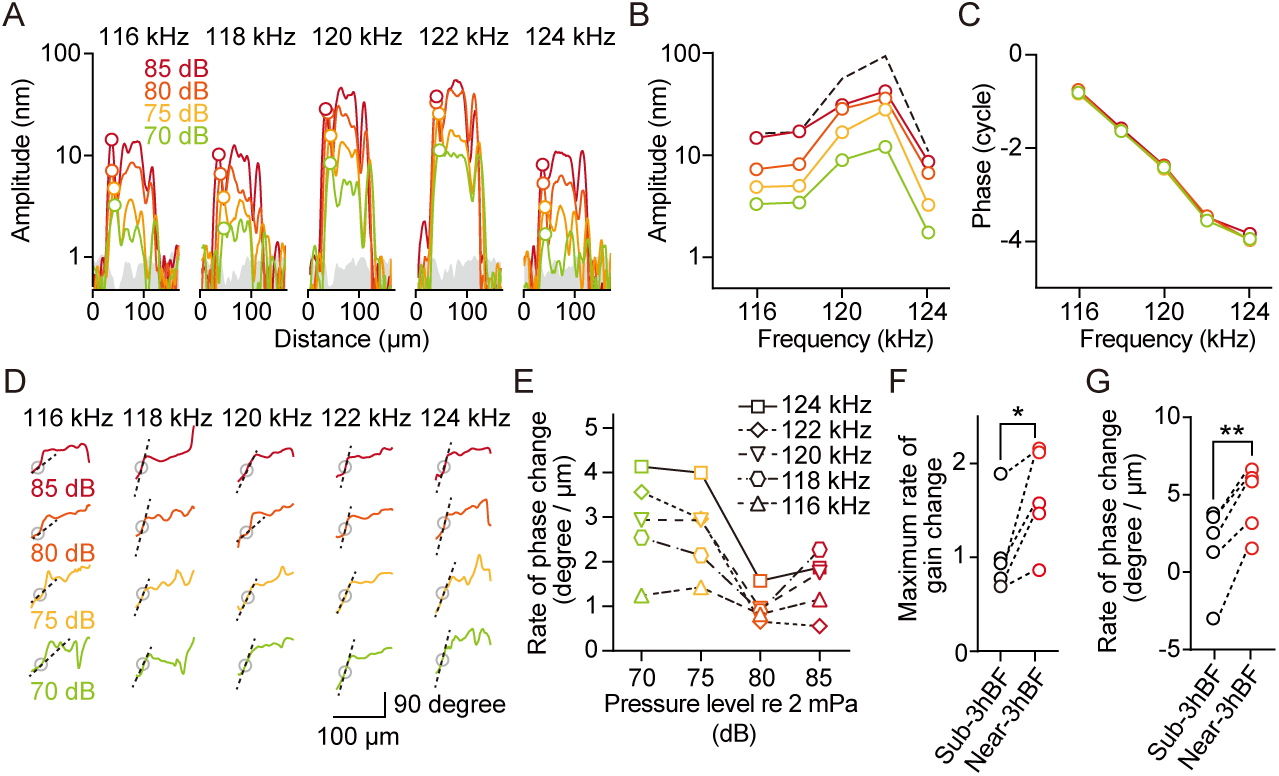
Optical coherence tomography (OCT) vibrometry analyses of the organ of Corti in the cochlear hook region during ultrasound stimulation beyond the hearing range. (A) Vibration amplitudes in response to stimulation from 116 kHz to 124 kHz with intensities of 70, 75, 80 and 85 dB. (B) Frequency distribution of the vibration amplitudes for the RL. The dashed black line shows the calculated linear increment in the amplitude for the 75 dB stimulus. The clear convergence of the peaks toward those under 85 dB stimulation indicates that the vibrations are compressive. (C) Frequency distribution of the phase for the RL in reference to the phase of the stimulation. (D) Relative phases in OHCs. For all stimuli, the phase at the RL significantly led that at the OHC body. Black open circles are the same points as the RL amplitudes, and dashed black lines indicate the rate of phase change near the RL. The rates of phase change were calculated from two pixels neighboring the RL. (E) Rates of phase change near the RL as a function of intensity. Different symbols indicate different frequencies. (F) Grouped maximum rate of gain in the near- and sub 3hBF groups (n = 5). Maximum rates of gain were collected from the gains between the different intensities by 10 dB with 10 dB increments. (G) Grouped rate of phase change near the RL in the sub- and near-3rdBF. * and **: significant differences (*p* < 0.32, and *p* < 0.05)

## Discussion

According to the conventional understanding of cochlear tonotopy, high-frequency sounds near the upper limit of the hearing range elicit a peak wave of epithelial motion in the hook region (Cooper & Rhode, 1992; Gelfand, 2010; Liberman, 1982; Manley, 2017; Robles & Ruggero, 2001; Ruggero & Temchin, 2002; Ulfendahl, 1997). How can the sensory epithelium resonate and transduce ultrasounds well above the animal’s hearing range? Previous studies on the sensory epithelium provide useful insights into our results. It has been reported that harmonic distortions of stimulus frequency occur in the apical and basal turns (Cooper, 1998; Dewey et al., 2021; Olson, 2004). Harmonic distortions result from hair cell nonlinear responses and/or from passive nonlinearity in the epithelial mechanics (Dewey et al., 2021). Our data are consistent with these reports; the frequency analysis of the hook region vibration exhibited a harmonic series in single tone stimulations (**Figure 4G**). Although CM include no harmonics in the single tone stimulations through the ossicle (**Figure S2**) (Dewey et al., 2021), hair cells transduces the stimulations that match not only the optimal frequency but also the harmonic frequencies into CM (**Figure 3** and **4**). This phenomenon is the principle of the CM synchronized with ultrasound beyond the hearing range, and plays a crucial role in both ultrasound-induced neuromodulation and ultrasonic hearing (Gavreau, 1948; Guo et al., 2018). In these situations, hair cells in the hook region transduce the ultrasound, but primarily cover the frequency near the upper limit of the hearing range. Consequently, afferent nerves connected to these hair cells convey information not about ultrasounds beyond the hearing range, but rather about audible sound. In ultrasonic hearing, although humans never perceive ultrasound through air conduction, we can hear it as high-frequency sound (Haeff & Knox, 1963). Furthermore, its frequency discrimination is less accurate than that within the hearing range (Yamashita, Nishimura, Nakagawa, Sakaguchi, & Hosoi, 2008). In ultrasound-induced neuromodulation, ultrasound produces extensive brain activation through the cochlear pathway (Guo et al., 2018). This study suggests a potential explanation for the occurrence of these sensations.

Our findings suggest that the cochlear compressive nonlinearity observed in ultrasound detection rests upon the MET current through OHCs. An important biomechanical function of OHCs that underlies the active process in mammalian cochleae is somatic electromotility: the cell bodies of OHCs change in length when the membrane potential is altered by the current through the MET channels on the hair bundles (Ashmore, 2008; Brownell, Bader, Bertrand, & de Ribaupierre, 1985). This modulation is produced by the membrane protein prestin (Futamata et al., 2022; Zheng et al., 2000). Although it has been reported that the motility of in OHCs is low-pass filtered (Gale & Ashmore, 1997; Santos-Sacchi, Iwasa, & Tan, 2019; Vavakou, Cooper, & van der Heijden, 2019), theoretical and experimental analyses of OHCs and prestin have suggested that the power output of somatic motility peaks at ultrasonic frequencies (Dewey et al., 2021; Rabbitt, 2020; Santos-Sacchi, Bai, & Navaratnam, 2023). OHCs display another mechanical function known as active hair bundle motility: the hair bundles of hair cells can oscillate spontaneously without external force and perform mechanical work to amplify inputs (Fettiplace, 2017; Hudspeth, 2008). To understand the interplay between somatic and bundle motilities in the hook region, further experiments, such as pharmacological perturbation of these motilities under in vivo and in vitro physiological conditions, may be required.

Tuning curves are not uniform at any level in the auditory pathway (Oertel, 2013). However, a tonotopic arrangement is known to be roughly conserved from the cochlear turns to the auditory cortex, and matches the hearing range in all mammals. In the classical model of cochlear frequency tuning for detecting a single tone, hair cells at a particular position are activated by a traveling wave, and send signals to afferent nerves (**Figure 6A**) (Gelfand, 2010; Hudspeth, 2013). In this study, we showed that hair cells in the hook region are electrically and mechanically resonant to both fundamental ultrasonic stimuli and their harmonics beyond the conventional range of hearing (**Figure 6B**). From the viewpoints of comparative and physiological hearing, interesting proposals have been put forward to explain sound detection in animals. First, many mammals can potentially sense ultrasound. In fact, recent studies have reported that some pinnipeds can hear ultrasound underwater (Cunningham & Reichmuth, 2016) and some fishes and frogs can perceive ultrasound in an attempt to avoid predation by echolocating animals or to facilitate communication (Feng et al., 2006; Wilson, Montie, Mann, & Mann, 2009). Ultrasonic hearing based on the harmonics detection by hair cells might be an evolutionary remnant conserved from these ancestral species, and traditional boundaries of the hearing range should be expanded to include ultrasound. Second, hair cells in other turns could be originally resonant to harmonics of the optimal frequency. This indicates that hair cells spaced at geometric intervals cooperatively function in moderate or strong single tone detection (**Figure 6C**). Additional physiological research on hair cells, the sensory epithelium, cochlear afferent nerves, and the auditory cortex is necessary to unravel the intricate mechanisms underlying harmonics detection in ultrasonic hearing and the collaborative role of hair cells in overall auditory function. These speculations provide a new principle for the treatment of deafness. For cochlear implants, otolaryngologists insert through the round window an electrode array that stimulates nerves and creates an auditory sensation (Lenarz, 2017). However, based on the theory of cochlear tonotopy, the electrodes in a limited position are active for each sound frequency. If the electrode array stimulates nerves in multiple positions, more electrical channels might be able to participate in improving patients’ discrimination of words or sounds. In addition, the relationship between ultrasonic hearing and deafness, such as acoustic trauma, ototoxicity, presbycusis, and Meniere’s disease, has not been fully studied (Lin, Thorne, Housley, & Vlajkovic, 2018). If hearing thresholds in ultrasonic hearing can capture the pre-symptomatic states of hair cells’ nonlinearity in these diseases, they could be used for prevention, prognostic prediction, and the development of treatments.

**Figure 6.**
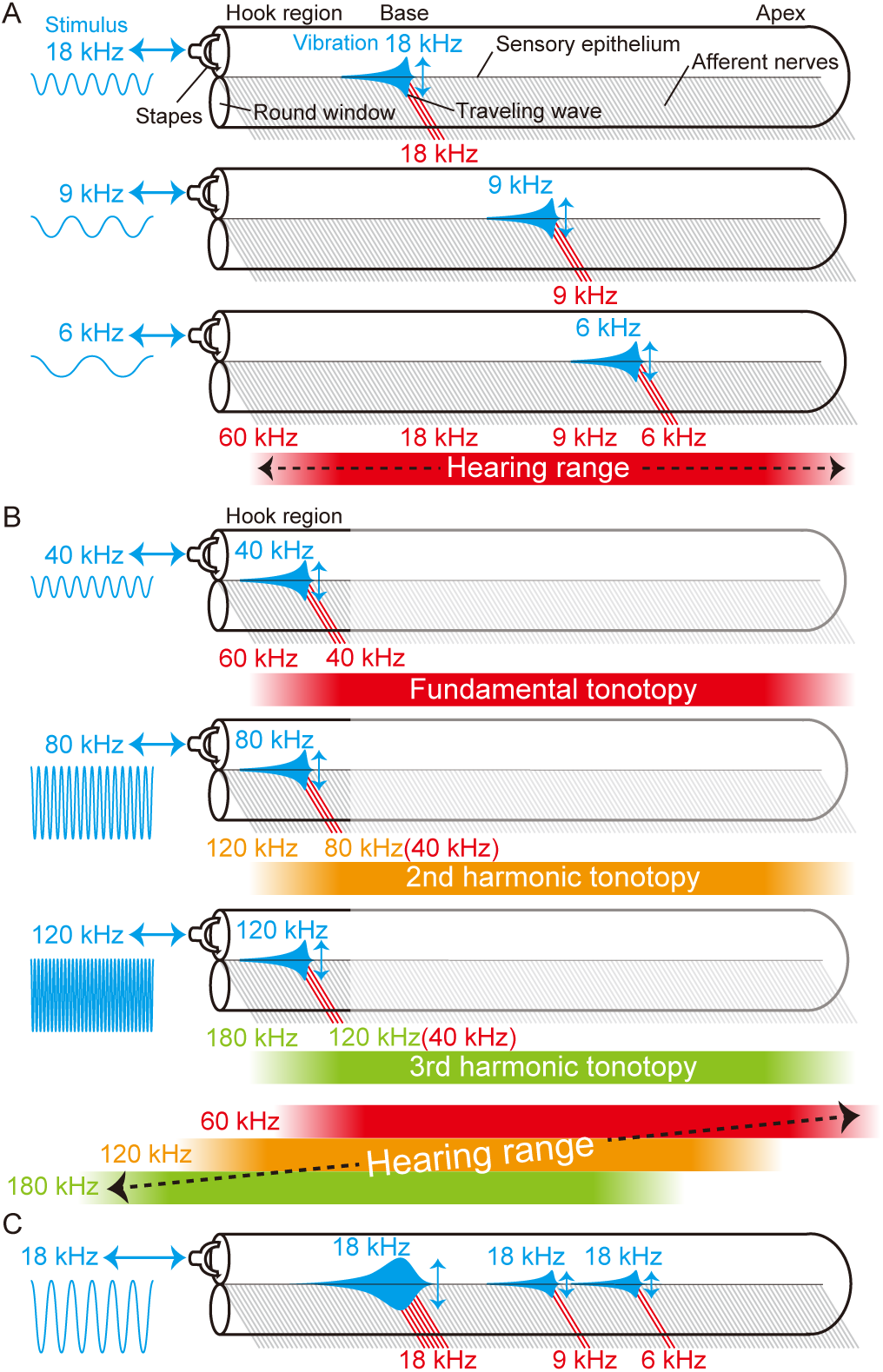
Depiction of stimulus-elicited vibrations on the sensory epithelium in the cochlea. (A) Basic concept of cochlear tonotopy. Each frequency of stimulation excites vibrations at a particular position. In this and subsequent illustrations, stimulations and vibrations are highlighted in blue. The red solid lines indicate activated afferent nerves under stimulation. The red shaded band in the lowest panel shows that the hearing range corresponds to the frequency distribution of the elicited vibrations and nerves. (B) Summary of our study. The upper, middle, and lower panels show the vibrations under stimulation at 40 kHz, 80 kHz, and 120 kHz, respectively. The red, orange and green shaded bands under the cochlea represent the frequency distributions of the fundamental, 2nd harmonic, and 3rd harmonic tonotopies, respectively. Although the applied frequencies are different, vibrations are evoked at the same position in the hook region. In the lowest panel, the sum of the frequency bands of the fundamental, 2nd harmonic, and 3rd harmonic tonotopies defines the hearing range. (C) Scheme of the predicted vibrations in a moderate or intense single audible tone stimulus. The frequency of the stimulus matches that of the vibrations on traveling waves, but evokes multiple traveling waves at different positions in subharmonic series of the fundamental frequency.

## Methods

### Animals and experimental procedures

The study included 190 healthy female guinea pigs weighing 200-400 g. The animals were treated in compliance with the Guiding Principles for the Care and Use of Animals in the field of Physiological Science set by the Physiological Society of Japan. The experimental protocol was approved by the Institutional Animal Research Committee of Gifu University (Permission Number: 2020-227). The experiments were carried out under the supervision of the committee and in accordance with the Guidelines for Animal Experiments of Gifu University and the Japanese Animal Protection and Management Law. Hartley guinea pigs (2-3 week of age; SLC Inc., Hamamatsu, Japan) were housed at the animal facility and kept under a 12-hr light and 12-hr dark cycle. All animal handling and reporting complied with the animal research: reporting of in vivo experiments (ARRIVE) guidelines.

For electrophysiological preparations, we first intramuscularly injected atropine sulfate (0.05 mg/kg) as a premedication. The animals were then anesthetized with an intraperitoneal injection of urethane (1.5 g/kg). After tracheostomy, the animals were artificially respirated with room air using a respirator (SN-408-7; Shinano Manufacturing, Japan). The cochlea was exposed using a method similar to that previously described for biophysical measurements (F. Nin et al., 2021; F. Nin, Reichenbach, Fisher, & Hudspeth, 2012; Ota et al., 2020). The head was fixed using a made-to-order stereotactic apparatus. The stapedial muscle and tensor tympani were surgically cut to avoid the effect of muscle reflexes during strong stimulation. Body temperature was monitored with a thermometer and maintained by a heating pad. Supplemental doses of anesthesia were administered to ensure areflexia to toe pinch. At the end of the experiment, anesthetized animals were injected intraperitoneally with an overdose of urethane.

For optical recordings, we performed artificial ventilation in urethane-anesthetized guinea pigs as described above. Using a ventrolateral approach, the cochlea was exposed by opening the bulla as previously described (F. Nin et al., 2021; F. Nin et al., 2012; Ota et al., 2020). An acrylic plate connected to a flexible base stage (SL20/M; Thorlabs, USA) was fixed to the head of the animal. The base stage and the body of the animal were placed on a movable platform under the OCT system. By orienting the head, the laser was focused on the hook region through the round-window membrane. Body temperature was monitored and maintained as described above.

Of the 190 animals included in this study, 128 were subjected to ABR and CM and 62 were subjected to OCT imaging and vibration recording. Without any perturbations, we measured the active ABR in more than 75% of the preparations. We recorded active CMs and epithelial vibrations in 31 and 15, respectively. During these recordings, we monitored the ABR thresholds before and after the experiment. If the ABR thresholds changed by more than 30 dB at the four frequencies of 16 kHz, 22 kHz, 30 kHz, and 40 kHz, we eliminated the data from the analysis. The low success rate of CMs and vibration recordings stems from invasive surgery, which often increases an animal’s hearing thresholds (F. Nin et al., 2021; F. Nin et al., 2012). Accordingly, we analyzed a limited number of samples that met the ABR threshold criteria. Nevertheless, the number of samples was almost equivalent to that in previous studies (F. Nin et al., 2021; F. Nin et al., 2012; Ota et al., 2020; Strimbu & Olson, 2022).

### Calibration of sound and ultrasound stimuli

We used a housed speaker (FT17H; Fostex, Japan) and a cuboidal piezoelectric actuator (PC4QM; Thorlabs, US) as stimulators. The speaker and actuator were powered using a function generator (WF1948; NF Corporation, Japan). For air-conducted sound stimulation, the tip of the speaker was inserted into the external ear canal. In contrast, for bone-conducted stimulation, the piezoelectric actuator was attached to the temporal bone using a ceramic rod glued on the actuator. Additionally, for ossicle stimulation, we used a tapered stainless rod and attached its tip to the malleus-incus complex. To avoid contamination by air-conducted sounds, the tympanic membrane was surgically removed during the bone conduction and ossicle stimulation experiments.

To calibrate the pressure of the air- and bone-conducted and ossicle stimulations, an ultrasound microphone (Sokolich ultrasonic probe microphone system, US) and a high sensitivity hydrophone (TC4034, Teledyne Marine REASON, US) with an amplifier (EC6081 mk2, Teledyne Marine REASON, US) were used. The microphone was connected to the tip of the speaker by a polyethylene tube for closed field sound stimulation, whereas the hydrophone was directly attached to the tips of the ceramic and stainless rods for bone-conducted and ossicle stimulations, respectively.

The intensity of a vibration stimulus unavoidably depends on the compression force applied to an object. To maintain the force constant, we monitored the force between the stimulator and the animal using a simple spring scale. During calibration, when the force was less than 2 N, the delivered pressure level decreased by more than 5 dB (**Figure S5A)**. Therefore, we maintained the force at 5 N in temporal bone stimulation. In ossicle stimulation, the compression force was reduced to 0.05 N because the surface area of the tip of the ceramic rod was approximately 100 times larger than that of the tip of the stainless rod (**Figure S5B)**.

### Electrophysiological measurements

The ABR was measured as described previously (Abaamrane et al., 2009). Stainless-steel needle electrodes were subcutaneously inserted in the parietal region under the pinnae and in the posterior region of the neck. For stimuli, 6 ms tone bursts with 0.5 ms rising and falling phases were generated by a function generator. The applied frequencies were 10, 16, 22, 30, 40, and 80 kHz in air-conducted stimulation, and 10, 16, 22, 30, 40, 80, 103, 127, 152, 176, 201, and 251 kHz in bone-conducted and ossicle stimulation. A piezoelectric actuator with ceramic and stainless rods did not emit secondary tone bursts during the stimulation (**Figure S6**). Individual signals emitted from the afferent auditory pathway were amplified 5000-fold and processed with an analog bandpass filter (300 Hz - 1 kHz) in a modified commercial amplifier (Model 3000, A-M Systems, US). These data were then digitally processed with a bandpass filter (300 Hz - 1 kHz), and 500 sweeps were averaged using LabVIEW (LabVIEW 2019 SP2; National Instruments, US). ABR thresholds were defined as the lowest pressure level at which wave III was detected.

The CM is a local field potential elicited by sound or vibration stimulation. The basic method of CM recording was similar to that used in previous studies (Honrubia & Ward, 1968; Tasaki, 1952). A made-to-order silver wire electrode and stainless-steel electrode were placed on the cochlear bone wall and parietal region under the pinnae, respectively. For the stimuli, 240 ms tone bursts of 80, 103, 127, 152, 176, 201, and 251 kHz with 20 ms rising and falling phases were generated by the function generator. Local field potentials were amplified 5000-fold and processed using an analog high-pass filter (1 kHz) in the amplifier. Acquired data were recorded by a digitizer with a sampling frequency of 2.4 MHz (PCIe-6374; National Instruments, US) and eight sweeps were averaged. We modified the commercial AC/DC amplifier that was originally utilized for recording neuronal signals. An analog lowpass filter was manually removed to record a high frequency voltage of more than 20 kHz. However, owing to the high input resistance of 10^15^ Ω, high-frequency signals greater than 1 kHz nonetheless deteriorated. Therefore, we obtained a voltage response curve as a function of stimulus frequency, and then corrected the acquired voltages using the values shown in **Figure S7**. Furthermore, because an actuator elicits electrical artifacts synchronized with the applied voltage frequency during stimulation, we defined the average amplitude of the recorded artifact as the LOD in the CM recording.

### Optical coherence tomography and vibrometry

We used a customized OCT system based on a commercial SD-OCT (Ganymede GAN621; Thorlabs, US) and a supercontinuum (SC) light source (SuperK FIU-15; NKT Photonics, Denmark). This modification was similar to that used in a previous OCT system termed the SCSD-OCT system (F. Nin et al., 2021), in which the light source was connected to an optical bandpass filter (SuperK SPLIT; NKT Photonics, Denmark), a fiber delivery system (SuperK CONNECT; NKT Photonics, Denmark), and a single-mode broadband fiber (FD7; NKT Photonics, Denmark). An objective lens (M Plan Apo NIR 20×, Mitutoyo, Japan) with a focal length of 10 mm and a depth of focus of 1.7 μm was used. The light power applied to the sample was 38.2 mW. The effective bandwidth and central wavelength of the spectrometer were approximately 300 nm and 900 nm, respectively. The axial pixel size in air was 0.56 μm. To analyze the anatomical properties of the sensory epithelium on OCT images, we estimated the refractive index of the cochlear lymph fluid to be 1.35. The sampling frequency of the OCT vibrometry was 248 kHz. The system therefore permitted direct measurement of vibrations at frequencies less than 124 kHz according to the sampling theorem. In the analysis of vibrations whose frequency exceeded 124 kHz, an alias signal for the stimulus frequency was acquired as previously reported (Dewey et al., 2021). To precisely evaluate the phase of the ultrasound, we used trigger signals generated with a high timing accuracy digitizer (National Instruments, PCIe-6374). LOD was assessed using the noise floor (NF) and standard deviation (SD) as LOD = NF + SD. To calculate the NF and SD, we used 30,000 scans of the vibration amplitudes at frequencies ranging from *f*-1500 Hz to *f*-500 Hz, where *f* is the stimulation frequency.

### Statistics and reproducibility

The means ± SDs are presented as descriptive statistics in **Figure 1, 3, S1**, and **S3**. CMs recorded in bone conduction and ossicle stimulation were compared using two-way ANOVA with interaction models for the two conditions and all the intensities studied in **Figure 3** and **S3**. The maximum rates of gain change and rates of phase change in the near-3hBF and sub-3hBF stimulation groups were compared using paired *t*-tests as shown in **Figure 5**. The *p* values were subjected to Bonferroni correction for multiple comparisons. All *n* values are indicated in the main text and the figures. All the statistical analyses were carried out using GraphPad Prism 9 (GraphPad Software, Inc., US)

## Data availability

All the data supporting the findings of this study are available within the paper, extended data figures, and figshare (10.6084/m9.figshare.25195850).

## Acknowledgments

We are especially grateful to Mr. Kazuo Tsukahara, and Drs. Takeru Ota, Shingo Murakami, and Yuuki Horii for their technical comments, Drs. A James Hudspeth, Tobias Reichenbach, Jonathan AN Fisher, and Francesco Gianoli for their critical reading of this manuscript, and Mr. Samuel Asare and Editage (www.editage.jp) for English language editing. This study was supported by the Toray Science Foundation 20-6109 (to F.N.), Shimadzu Science Foundation 2020-13 (to F.N.), UBE Foundation 2020-55 (to F.N.), Nakatani Foundation for Advancement of Measuring Technologies in Biomedical Engineering 2021K013, 2022S107 (to F.N. and K.H.), The Salt Science Research Foundation 2228 (to F.N.), Suzuken Memorial Foundation 23-022 (to F.N.), The Naito Foundation (to F.N.), The Uehara Memorial Foundation 202120167, 202110177 (to F.N. and K.H.), Seiko Instruments Advanced Technology Foundation Research Grants 2023 RGR0503 (to K.H.), Ogawa Memorial Foundation (to K.H.), Kowa Life Science Foundation 2022-15 (to K.H.), and Grant-in-Aid for Young Scientists 22K15364 (to K.H.), Scientific Research (B) 23H03307 (to C.A.), Challenging Exploratory Research 22K19617 (to C.A.), and Fostering Joint International Research (A) 19KK0364 (to S.C.) from the Ministry of Education, Culture, Sports, Science and Technology of Japan.

## Author Contributions

F. N. designed the research; K. H., B. O., N. N., I. M., and F. N. performed the research; S. C., T. O., and F. N. contributed new reagents/analytic tools; K. H., B. O., C. A., and F. N. analyzed the data; and K.H. and N. wrote the paper. K. H. and B. O. contributed equally to this work;

## References

Abaamrane, L., Raffin, F., Gal, M., Avan, P., & Sendowski, I. (2009). Long-term administration of magnesium after acoustic trauma caused by gunshot noise in guinea pigs. Hear Res, 247(2), 137–145. doi:10.1016/j.heares.2008.11.005

Ashmore, J. (2008). Cochlear outer hair cell motility. Physiol Rev, 88(1), 173–210. doi:10.1152/physrev.00044.2006

Bekesy, G. v. (1960). Experiments in hearing (E. G. Wever Ed.): McGraw-Hill Book Company.

Brownell, W. E., Bader, C. R., Bertrand, D., & de Ribaupierre, Y. (1985). Evoked mechanical responses of isolated cochlear outer hair cells. Science, 227(4683), 194–196. doi:10.1126/science.3966153

Cooper, N. P. (1998). Harmonic distortion on the basilar membrane in the basal turn of the guinea-pig cochlea. J Physiol, 509 ( *Pt 1*)(Pt 1), 277–288. doi:10.1111/j.1469-7793.1998.277bo.x

Cooper, N. P., & Rhode, W. S. (1992). Basilar membrane mechanics in the hook region of cat and guinea-pig cochleae: sharp tuning and nonlinearity in the absence of baseline position shifts. Hear Res, 63(1-2), 163–190. doi:10.1016/0378-5955(92)90083-y

Corso, J. (1963). Bone-conduction thresholds for sonic and ultrasonic frequencies. J Acoust Soc Am, 35, 1738–1743.

Cunningham, K. A., & Reichmuth, C. (2016). High-frequency hearing in seals and sea lions. Hear Res, 331, 83–91. doi:10.1016/j.heares.2015.10.002

Dewey, J. B., Altoe, A., Shera, C. A., Applegate, B. E., & Oghalai, J. S. (2021). Cochlear outer hair cell electromotility enhances organ of Corti motion on a cycle-by-cycle basis at high frequencies in vivo. Proc Natl Acad Sci U S A, 118(43). doi:10.1073/pnas.2025206118

Dieroff, H. G., & Ertel, H. (1975). Some thoughts on the perception of ultrasonics by man. Arch Otorhinolaryngol, 209(4), 277–290. doi:10.1007/BF00456548

Feng, A. S., Narins, P. M., Xu, C. H., Lin, W. Y., Yu, Z. L., Qiu, Q.,… Shen, J. X. (2006). Ultrasonic communication in frogs. Nature, 440(7082), 333–336. doi:10.1038/nature04416

Fettiplace, R. (2017). Hair cell transduction, tuning, and synaptic transmission in the mammalian cochlea. Compr Physiol, 7(4), 1197–1227. doi:10.1002/cphy.c160049

Fisher, J. A. N., & Gumenchuk, I. (2018). Low-intensity focused ultrasound alters the latency and spatial patterns of sensory-evoked cortical responses in vivo. J Neural Eng, 15(3), 035004. doi:10.1088/1741-2552/aaaee1

Futamata, H., Fukuda, M., Umeda, R., Yamashita, K., Tomita, A., Takahashi, S.,… Nureki, O. (2022). Cryo-EM structures of thermostabilized prestin provide mechanistic insights underlying outer hair cell electromotility. Nat Commun, 13(1), 6208. doi:10.1038/s41467-022-34017-x

Gale, J. E., & Ashmore, J. F. (1997). An intrinsic frequency limit to the cochlear amplifier. Nature, 389(6646), 63–66. doi:10.1038/37968

Gavreau, V. (1948). Audibillite de sons de frequence elevee. Compt Rendu, 226, 2053–2054.

Gelfand, S. A. (2010). Cochlear mechanism and process (5th edition ed.).

Guo, H., Hamilton Ii, M., Offutt, S. J., Gloeckner, C. D., Li, T., Kim, Y.,… Lim, H. H. (2018). Ultrasound produces extensive brain activation via a cochlear pathway. Neuron, 99(4), 866. doi:10.1016/j.neuron.2018.07.049

Haeff, A. V., & Knox, C. (1963). Perception of ultrasound. Science, 139(3555), 590–592. doi:10.1126/science.139.3555.590

Hoffman, B. U., Baba, Y., Lee, S. A., Tong, C. K., Konofagou, E. E., & Lumpkin, E. A. (2022). Focused ultrasound excites action potentials in mammalian peripheral neurons in part through the mechanically gated ion channel PIEZO2. Proc Natl Acad Sci U S A, 119(21), e2115821119. doi:10.1073/pnas.2115821119

Honrubia, V., & Ward, P. H. (1968). Longitudinal distribution of the cochlear microphonics inside the cochlear duct (guinea pig). J Acoust Soc Am, 44(4), 951–958. doi:10.1121/1.1911234

Hosoi, H., Imaizumi, S., Sakaguchi, T., Tonoike, M., & Murata, K. (1998). Activation of the auditory cortex by ultrasound. Lancet, 351(9101), 496–497. doi:10.1016/S0140-6736(05)78683-9

Hudspeth, A. J. (2008). Making an effort to listen: mechanical amplification in the ear. Neuron, 59(4), 530–545. doi:10.1016/j.neuron.2008.07.012

Hudspeth, A. J. (2013). Principles of neural science: The inner ear (Fifth edition ed.): Mc Graw Hill.

Koizumi, T., Nishimura, T., Yamashita, A., Yamanaka, T., Imamura, T., & Hosoi, H. (2014). Residual inhibition of tinnitus induced by 30-kHz bone-conducted ultrasound. Hear Res, 310, 48–53. doi:10.1016/j.heares.2014.01.011

Kubanek, J., Shi, J., Marsh, J., Chen, D., Deng, C., & Cui, J. (2016). Ultrasound modulates ion channel currents. Sci Rep, 6, 24170. doi:10.1038/srep24170

Lenarz, T. (2017). Cochlear implant - state of the art. GMS Curr Top Otorhinolaryngol Head Neck Surg, 16, Doc04. doi:10.3205/cto000143

Liberman, M. C. (1982). The cochlear frequency map for the cat: labeling auditory-nerve fibers of known characteristic frequency. J Acoust Soc Am, 72(5), 1441–1449. doi:10.1121/1.388677

Lin, S. C. Y., Thorne, P. R., Housley, G. D., & Vlajkovic, S. M. (2018). Resistance to neomycin ototoxicity in the extreme basal (hook) region of the mouse cochlea. Histochem Cell Biol, 150(3), 281–289. doi:10.1007/s00418-018-1683-8

Manley, G. A. (2017). The cochlea: What It Is, where Is came from, and what Is special about it.

Nin, F., Choi, S., Ota, T., Qi, Z., & Hibino, H. (2021). Optimization of spectral-domain optical coherence tomography with a supercontinuum source for in vivo motion detection of low reflective outer hair cells in guinea pig cochleae. Opt Rev, 28, 239–254.

Nin, F., Reichenbach, T., Fisher, J. A., & Hudspeth, A. J. (2012). Contribution of active hair-bundle motility to nonlinear amplification in the mammalian cochlea. Proc Natl Acad Sci U S A, 109(51), 21076–21080. doi:10.1073/pnas.1219379110

Nishimura, T., Nakagawa, S., Sakaguchi, T., & Hosoi, H. (2003). Ultrasonic masker clarifies ultrasonic perception in man. Hear Res, 175(1-2), 171–177. doi:10.1016/s0378-5955(02)00735-9

Nishimura, T., Okayasu, T., Uratani, Y., Fukuda, F., Saito, O., & Hosoi, H. (2011). Peripheral perception mechanism of ultrasonic hearing. Hear Res, 277(1-2), 176–183. doi:10.1016/j.heares.2011.01.004

Nitsche, M. A., Cohen, L. G., Wassermann, E. M., Priori, A., Lang, N., Antal, A.,… Pascual-Leone, A. (2008). Transcranial direct current stimulation: State of the art 2008. Brain Stimul, 1(3), 206–223. doi:10.1016/j.brs.2008.06.004

Oertel, D. D., A. J. (2013). Principles of neural science: The auditory central nerve system (Fifth edition ed.).

Ohyama, K., Kusakari, J., & Kawamoto, K. (1985). Ultrasonic electrocochleography in guinea pig. Hear Res, 17(2), 143–151. doi:10.1016/0378-5955(85)90017-6

Olson, E. S. (2004). Harmonic distortion in intracochlear pressure and its analysis to explore the cochlear amplifier. J Acoust Soc Am, 115(3), 1230–1241. doi:10.1121/1.1645611

Ota, T., Nin, F., Choi, S., Muramatsu, S., Sawamura, S., Ogata, G.,… Hibino, H. (2020). Characterisation of the static offset in the travelling wave in the cochlear basal turn. Pflugers Arch, 472(5), 625–635. doi:10.1007/s00424-020-02373-6

Rabbitt, R. D. (2020). The cochlear outer hair cell speed paradox. Proc Natl Acad Sci U S A, 117(36), 21880–21888. doi:10.1073/pnas.2003838117

Reichenbach, T., & Hudspeth, A. J. (2010). Dual contribution to amplification in the mammalian inner ear. Phys Rev Lett, 105(11), 118102. doi:10.1103/PhysRevLett.105.118102

Robles, L., & Ruggero, M. A. (2001). Mechanics of the mammalian cochlea. Physiol Rev, 81(3), 1305–1352. doi:10.1152/physrev.2001.81.3.1305

Ruggero, M. A., & Temchin, A. N. (2002). The roles of the external, middle, and inner ears in determining the bandwidth of hearing. Proc Natl Acad Sci U S A, 99(20), 13206–13210. doi:10.1073/pnas.202492699

Santos-Sacchi, J., Bai, J. P., & Navaratnam, D. (2023). Megahertz sampling of prestin (SLC26a5) voltage-sensor charge movements in outer hair cell membranes reveals ultrasonic activity that may support electromotility and cochlear amplification. J Neurosci. doi:10.1523/JNEUROSCI.2033-22.2023

Santos-Sacchi, J., Iwasa, K. H., & Tan, W. (2019). Outer hair cell electromotility is low-pass filtered relative to the molecular conformational changes that produce nonlinear capacitance. J Gen Physiol, 151(12), 1369–1385. doi:10.1085/jgp.201812280

Sato, T., Shapiro, M. G., & Tsao, D. Y. (2018). Ultrasonic neuromodulation causes widespread cortical activation via an indirect auditory mechanism. Neuron, 98(5), 1031–1041 e1035. doi:10.1016/j.neuron.2018.05.009

Strimbu, C. E., & Olson, E. S. (2022). Salicylate-induced changes in organ of Corti vibrations. Hear Res, 423, 108389. doi:10.1016/j.heares.2021.108389

Tasaki, I. D., H.; Legouix, J. P. (1952). The space-time pattern of the cochlear microphonics (guinea pig), as recorded by differential electrodes. J Acoust Soc Am, 24(5), 502–519.

Tchumatchenko, T., & Reichenbach, T. (2014). A cochlear-bone wave can yield a hearing sensation as well as otoacoustic emission. Nat Commun, 5, 4160. doi:10.1038/ncomms5160

Tyler, W. J., Tufail, Y., Finsterwald, M., Tauchmann, M. L., Olson, E. J., & Majestic, C. (2008). Remote excitation of neuronal circuits using low-intensity, low-frequency ultrasound. PLoS One, 3(10), e3511. doi:10.1371/journal.pone.0003511

Ulfendahl, M. (1997). Mechanical responses of the mammalian cochlea. Prog Neurobiol, 53(3), 331–380. doi:10.1016/s0301-0082(97)00040-3

Vavakou, A., Cooper, N. P., & van der Heijden, M. (2019). The frequency limit of outer hair cell motility measured in vivo. Elife, 8. doi:10.7554/eLife.47667

Wever, E. G., & Bray, C. W. (1930). Action currents in the auditory nerve in response to acoustical stimulation. Proc Natl Acad Sci U S A, 16(5), 344–350. doi:10.1073/pnas.16.5.344

Wilson, M., Montie, E. W., Mann, K. A., & Mann, D. A. (2009). Ultrasound detection in the Gulf menhaden requires gas-filled bullae and an intact lateral line. J Exp Biol, 212(Pt 21), 3422–3427. doi:10.1242/jeb.033340

Yamashita, A., Nishimura, T., Nakagawa, S., Sakaguchi, T., & Hosoi, H. (2008). Assessment of ability to discriminate frequency of bone-conducted ultrasound by mismatch fields. Neurosci Lett, 438(2), 260–262. doi:10.1016/j.neulet.2008.03.086

Zheng, J., Shen, W., He, D. Z., Long, K. B., Madison, L. D., & Dallos, P. (2000). Prestin is the motor protein of cochlear outer hair cells. Nature, 405(6783), 149–155. doi:10.1038/35012009

