## Supplementary figures and images for "The cochlear hook region detects harmonics beyond the canonical hearing range"

### Figure S1

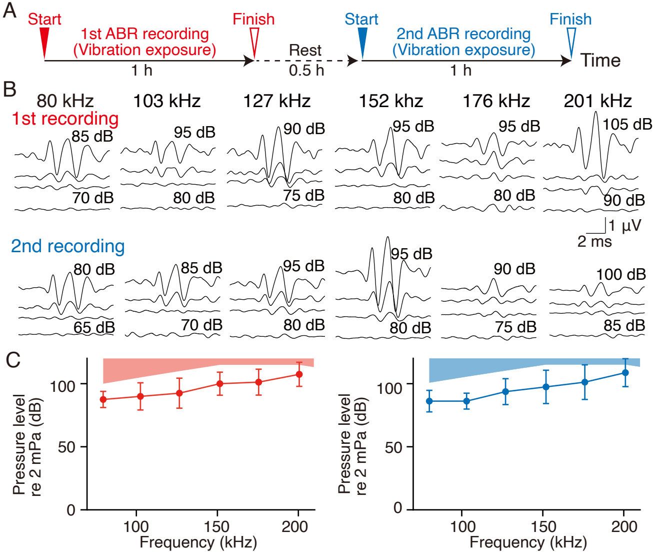

### Figure S2

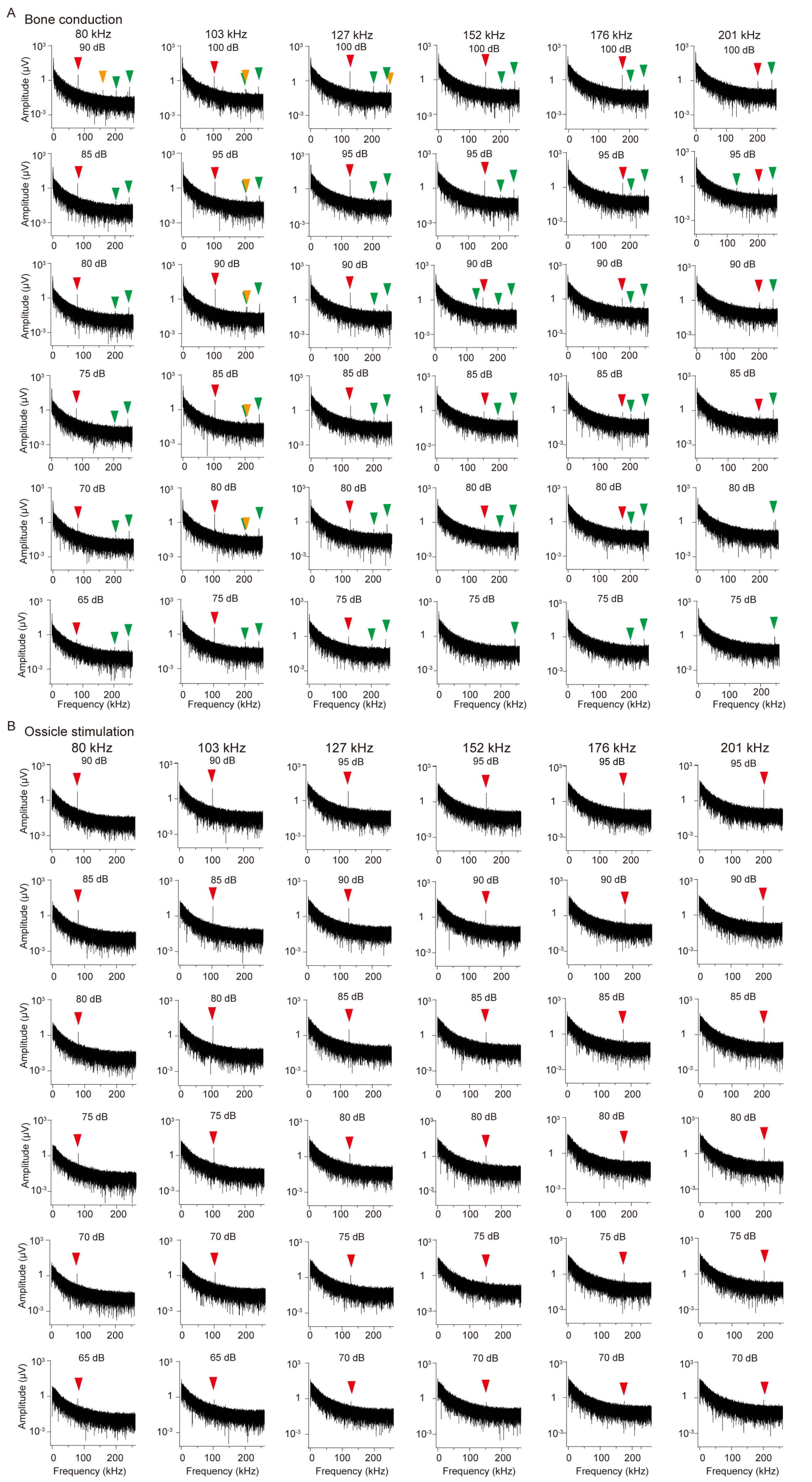

### Figure S3

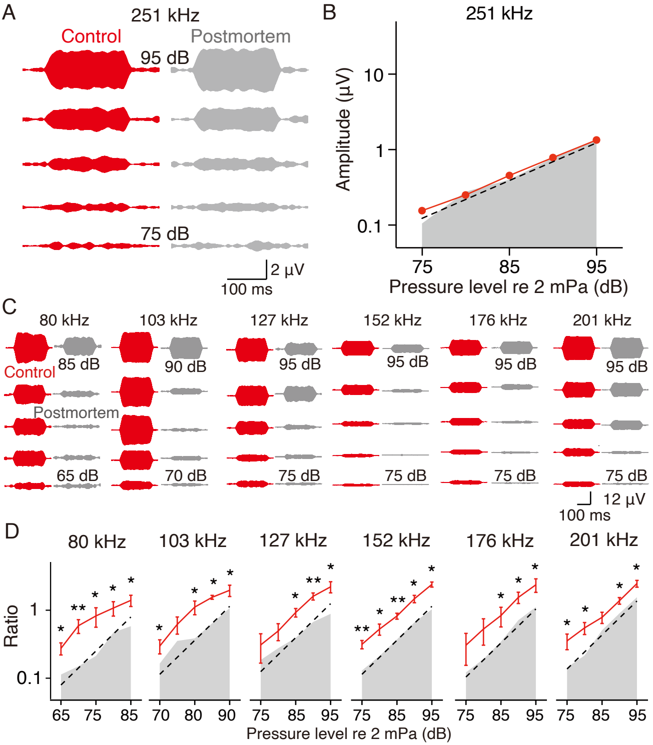

### Figure S4

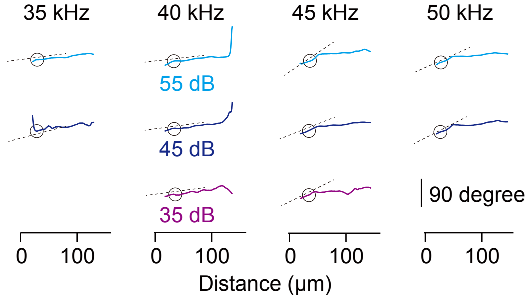

### Figure S5

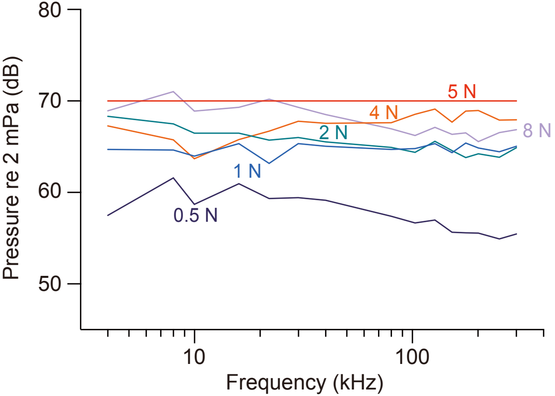

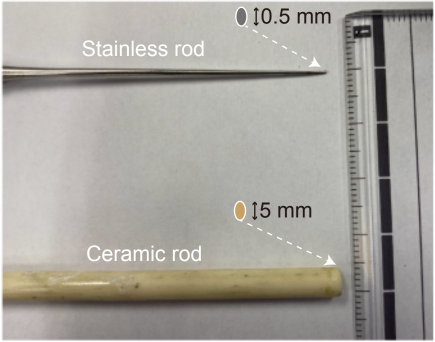

### Figure S6

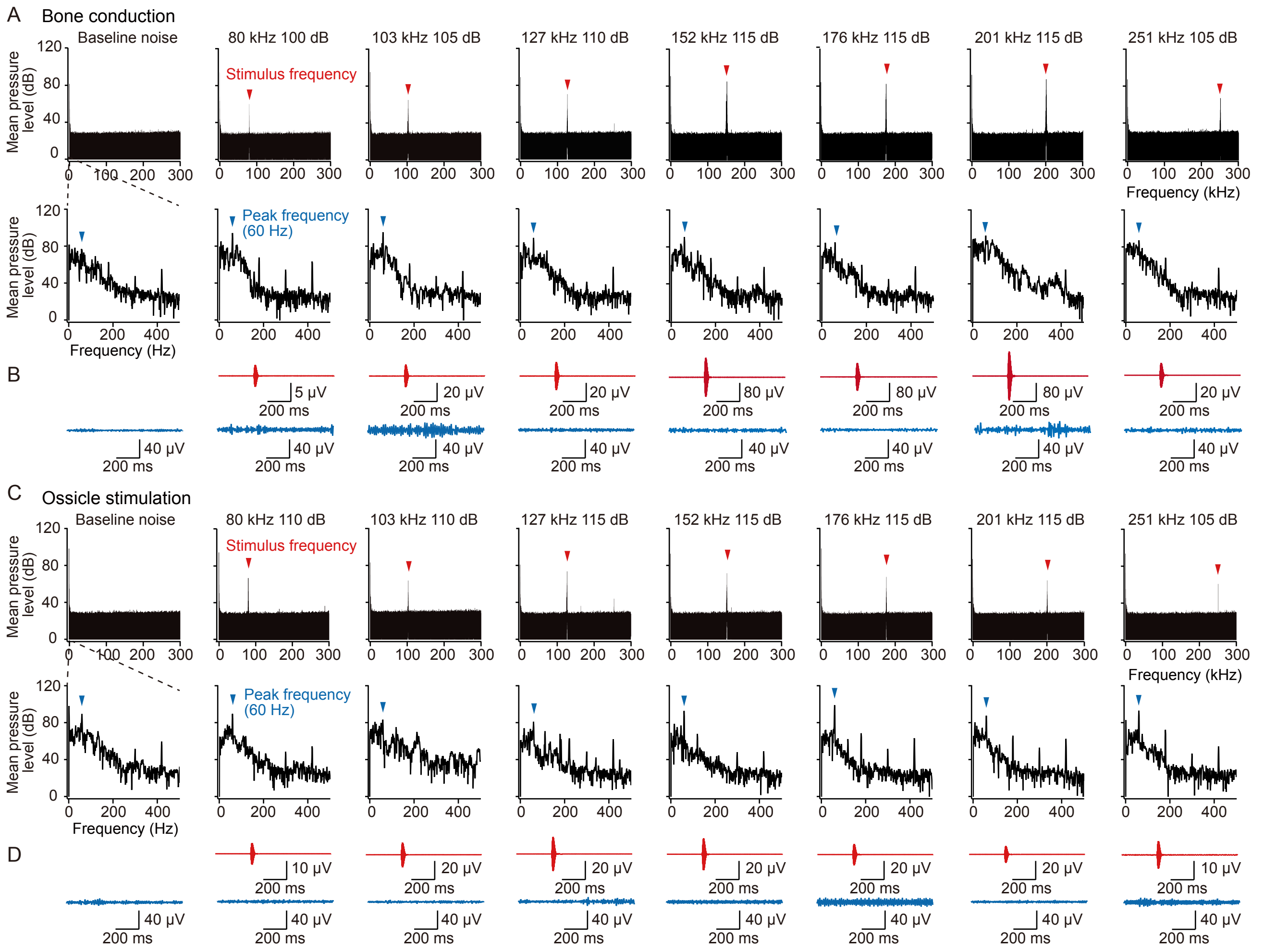

### Figure S7

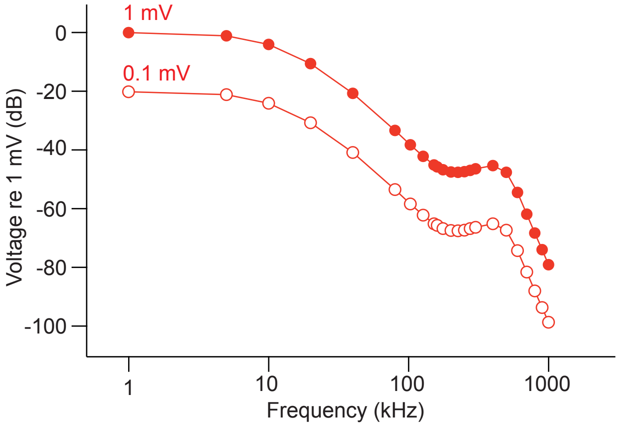
